# Transcriptomic response of an Antarctic yeast *Rhodotorula* sp. USM-PSY62 to temperature changes

**DOI:** 10.1101/2020.05.07.080796

**Authors:** Cleo-Nicole Chai, Hok-Chai Yam, Nurlina Rosli, Azali Azlan, Ghows Azzam, Mardani Abdul Halim, Nazalan Najimudin

**Author notes:** Corresponding authors: Nazalan Najimudin and Mardani Abdul Halim, Address: School of Biological Sciences, Universiti Sains Malaysia, 11800 Penang, Malaysia.

## Abstract

*Rhodotorula* sp. (USM-PSY62) is a psychrophilic yeast isolated from Antarctic sea ice and it grows optimally at 15 °C. This study was set up to observe how USM-PSY62 adapted to fluctuations in temperature. During cold adaptation, an elevated transcription of the CorA magnesium transporter gene in USM-PSY62 indicated a higher requirement for magnesium ions in order to gain additional enzyme cofactors or maintain cytoplasmic fluidity. The HepA homologue coding for DNA/RNA helicase was also over-expressed in cold condition possibly to reorganize secondary structures of DNA and RNA. An up-regulation of the catalase gene was also observed reflecting an increment in the concentration of reactive oxygen species and fluctuations in the associated antioxidant system. The *YOP1* gene, which encodes a membrane protein associated with protein transport and membrane traffic, was the most down-regulated under cold shock condition. The genes responsible for the structural maintenance of chromosome (*SMC*) were also down-regulated when the temperature was shifted to 0 °C. Upon cold shock, the gene for heat shock factor protein 1 (*HSF*1) was also down-regulated. Hsf1 is a transcriptional regulator which regulate the heat shock responses. Although USM-PSY62 showed some common adaptive strategies as in several other psychrophilic organisms, new mechanisms were also uncovered.

## Introduction

Antarctica is generally known as the coldest continent with an average annual temperature of −55° and wintry conditions that can go to as low as −85 °C (Stonehouse 2004). With an estimated area of 14 million square kilometres, the continent is larger than Australia or Europe with several unique environmental niches involving different types of soil, sediments, rocks, snow and deep ice (Russell 2006). Most of these settings are also vastly unexplored.

The continent is also a sanctuary for a lot of microorganisms with remarkable arrays of metabolisms (Lohan and Johnston 2005). These microorganisms represent the most abundant cold-adapted organisms on earth with high species diversity as well as biomass (Feller and Gerday 2003). These are known as psychrophiles due to their cold-adapting nature. Psychrophiles are distinct from psychrotrophs that can grow at the same range of low temperatures as the former, but their maximal growth temperature can be above 20 °C (Morita 1975). Both psychrophiles and psychrotrophs can be found in the same environment. However, the psychrotrophic organisms are usually superior in number in niches with temperatures that fluctuate above the psychrophilic range. Both types play significant roles in the biodegradation of organic compounds as well as the cycling of essential nutrients (Russell 1990; Vishniac 2006).

Yeasts are eukaryotes and many species of yeasts have unique physiological abilities that allow them to survive in a wide range of surroundings (Hagler 1987; Starmer and Lachance 2010). Various studies showed that yeasts can adapt well to cold environments (Margesin et al. 2003; Turkiewicz et al. 2003). The presence of psychrophilic and psychrotrophic yeasts in cold ecosystems have been well documented (Fell 2006, Raspor 2006, Vishniac 2006). The most common yeast species found in cold environments are from the basidiomycetous group, though some ascomycetous yeasts have also been previously described (Branda et al. 2010; Connell et al. 2008; de García et al. 2010). Examples of species of yeast isolated from cold environments are *Mrakia robertii, Mrakia blollopis, Mrakia niccombsii* (Thomas-Hall et al. 2010), *Rhodotorula arctica* (Vishniac and Takashima 2010), *Dioszegia antarctica* and *Diozegia cryoxerica* (Connell et al. 2010). These illustrate the richness and diversity in these extreme environments highlighting the opportunity to further explore their diversity.

The current period of global warming poses a lot of challenge to the microbial diversity in Antarctica and it is therefore essential to understand the associated effects of the temperature increment. It is crucial to find out how psychrophilic microbes adapt to fluctuations in their growth temperatures and shed new lights on their survival mechanisms (Deegenaars and Watson 1998). Psychrophiles grow and reproduce in low temperature environments. Among their survival methods include the ability to maintain functionality of enzymes that tend to rigidify when a descent in temperature occurs as well preserve the flexibility of the cytoplasmic membranes to perform signalling and biochemical reactions (Ratkowsky et al. 2005). Cold adaptive enzymes developed more flexibility at lower temperature (Hoyoux et al. 2004) and active sites became more accessible for enzymatic reactions to occur (Aghajari et al. 2003). The evolution of cold-shock proteins is another strategy to adapt to extreme low temperatures. These proteins are regularly synthesized in various organisms during extended period of growth in the cold (Pikuta et al. 2007). In this study, a transcriptomic analysis was performed on an Antarctic yeast, *Rhodotorula* sp. USM-PSY62, to bring to light more unique adaptive mechanisms to temperature fluctuations and understand how these eukaryotic extremophiles thrive in their environments. This will also allow us to understand how psychrophilic microbes respond to environmental changes in a fluctuating ecosystem.

## Materials and methods

### Yeast strain and maintenance

The basidiomycete strain *Rhodotorula* sp. USM-PSY62 was isolated from an Antarctic sea ice sample collected at Casey Station, Antarctica (Ong 2006). USM-PSY62 was grown and maintained in YPD medium (1% yeast extract, 2% polypeptone and 2% dextrose) at 15 °C (Vellanki 2013). For long term storage, a cell suspension was prepared in YPD supplemented with 20% glycerol and maintained frozen at −80 °C.

### Exposure of USM-PSY62 to different temperatures

USM-PSY62 was initially grown in YPD broth as a starter culture at its optimum growth temperature of 15°C for five days. A small volume of starter culture was then inoculated into 500 ml flasks containing 100 ml of fresh YPD broth to an OD_600nm_ of approximately 0.1. These flasks were incubated in a shaker at 150 rpm and 15°C for 7 days until mid-log phase was reached. Thereafter, individual cultures were then transferred to 0°C, 5°C, 15°C and 21°C for 24 hours of incubation shaken at 150 rpm at two replicates for each temperature. After the exposure period, the cells were harvested by centrifugation. The supernatant was discarded and the cell pellets were immediately snapped frozen in liquid nitrogen. They were then stored at - 70°C prior to RNA extraction.

### Total RNA extraction and RNA-Seq

The USM-PSY62 cells were homogenized using liquid nitrogen and a grinder and the total RNA was extracted using TRIzol® Reagent (Thermo Fisher Scientific, USA) as recommended by the manufacturer. DNA was removed using TURBO DNase (Thermo Fisher Scientific, USA). The integrity of the extracted RNA was assessed using denaturing agarose gel electrophoresis and RNA concentration was quantified using Nanodrop-1000 (Thermo Fisher Scientific, USA). The cDNA libraries from the purified RNA were sequenced using the Illumina HiSeq 2000 platform. The transcriptome results were validated through quantitative real time PCR (qRT-PCR).

### *de novo* transcriptome assembly and functional annotation

A *de novo* assembly process was performed on the clean data generated from the transcriptome sequencing. The assembler used in this study was Trinity version r2013-11-10 (Grabherr et al. 2011; Haas et al. 2013). The minimum assembled contig length was changed to 100 and the option for high expectation of paired reads with high density with UTR overlap was used (--jaccard_clip). The processed transcripts were annotated using Trinotate suite (http://trinotate.sourceforge.net/) which classified them using the eggNOG (evolutionary genealogy of genes: Non-supervised Orthologous Groups) public database of orthology relationships, gene evolutionary histories and functional annotations. To further refine the transcriptome assembly, transcripts having more than 60% BLASTN alignment match to bacterial species were filtered away. Additionally, transcripts were mapped to a few selected organisms, namely *Escherichia coli, Saccharomyces cerevisiae, Rhodotorula mucilaginosa* and *Caenorhabditis elegans*, to observe their similarities.

### Transcripts abundance estimation and differential expression analysis

Transcripts counts were performed by using the RNAseq Expectation-Maximization (RSEM) method (Li and Dewey 2011). The gene expression level was normalized as the number of fragments per kilobase of exon region per million mapped reads (FPKM). To estimate the differentially expressed transcripts for each pairwise comparison of the samples, EdgeR software was used (Robinson et al. 2010). Then, the Trimmed Mean of M-values (TMM) normalization between samples was performed to adjust for the different library sizes. Significance was established by comparing the gene expression levels which were normalized by TMM. The differentially expressed (DE) transcripts extracted were those that were differentially expressed by at least two-fold at a false discovery-corrected statistical significance set to at most 0.001.

## Results

### Raw data and sequence assembly

In general, the post filtering and assembly raw data generated from RNA-Seq was relatively uniform with an average of approximately 25 million reads per sample. The average GC value of all the samples was 61.52 %. The summary of raw data generated from RNA-Seq is shown in Table 1. The *de novo* assembly performed on the raw data generated assembled the reads into a total of 34170 contig sequences ranging from 101 bp to 9227 bp. The median and mean length of the contigs were 309 and 673 bp, respectively, with an N50 value of 1400 bp. The overall statistics generated by the Trinity *de novo* assembly is as presented in Table 2. The most abundant contig lengths in the assembly were in the range of 101 bp to 500 bp while the least were in the range of 2501 bp to 3000 bp as shown in Figure 1. Results from the alignment to selected species revealed that 2.99 % of the transcripts mapped to the genome of *Escherichia coli*, 0.17 % to *Saccharomyces cerevisiae*, 8.92 % to *Rhodotorula mucilaginosa* and 0.57 % to *Caenorhabditis elegans*. The raw data can be accessed in NCBI under BioProject ID PRJNA522043.

**Table 1.** RNA-Seq sequencing output statistics. Total clean reads were obtained after removing the adapters and low-quality sequences. An average of 25 million reads were obtained for each sample with a GC value of 61.52 %.

| **Sample** | **Total raw reads** | **Total clean reads** | **Total clean nucleotides** | **Q20 %** | **GC %** |
| --- | --- | --- | --- | --- | --- |
| **0°C_Rep 1** | 28,478,626 | 24,479,720 | 2,447,972,000 | 97.73 % | 61.25 % |
| **0°C_Rep 2** | 25,860,066 | 24,253,342 | 2,425,334,200 | 97.68 % | 61.43 % |
| **5°C_Rep 1** | 26,258,816 | 24,270,378 | 2,427,037,800 | 97.78 % | 61.15 % |
| **5°C_Rep 2** | 25,904,280 | 24,214,220 | 2,421,422,000 | 97.72 % | 61.47 % |
| **15°C_Rep 1** | 26,689,456 | 24,338,464 | 2,433,846,400 | 97.72 % | 61.40 % |
| **15°C_Rep 2** | 27,375,710 | 23,654,056 | 2,365,405,600 | 92.23 % | 61.68 % |
| **21°C_Rep 1** | 26,645,038 | 24,613,284 | 2,461,328,400 | 97.69 % | 61.98 % |
| **21°C_Rep 2** | 28,342,708 | 24,505,100 | 2,450,510,000 | 92.05 % | 61.81 % |

**Table 2.**
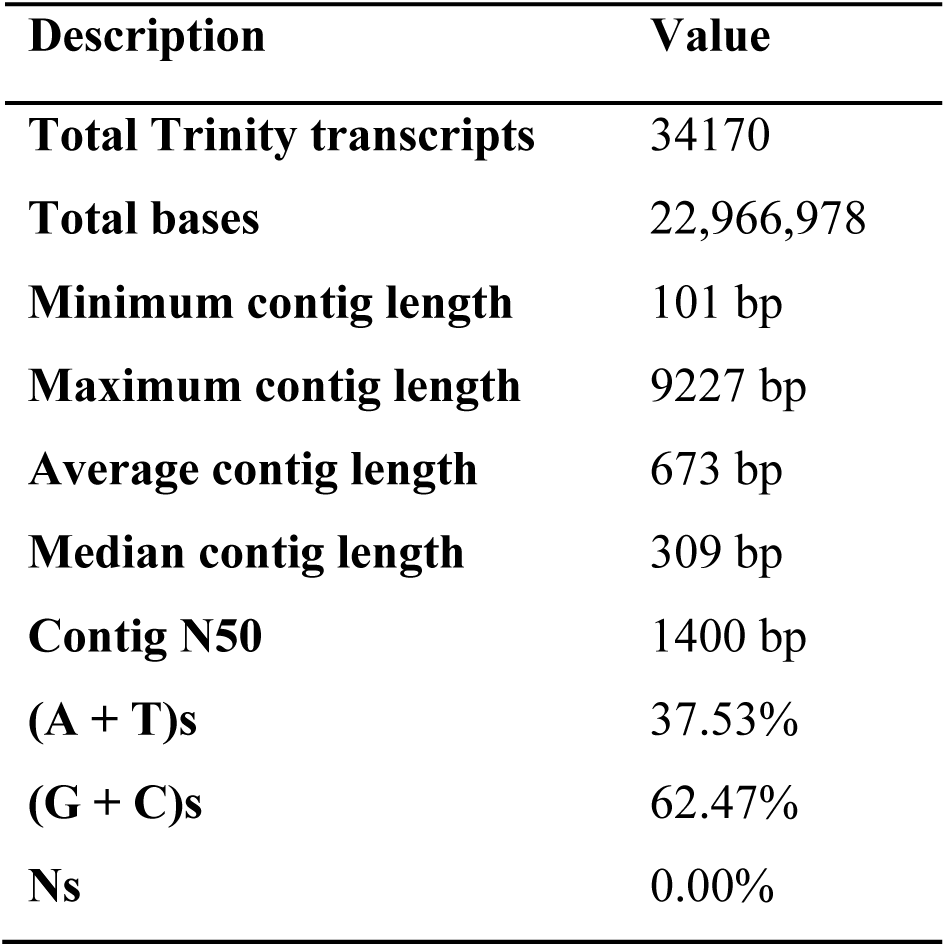
Trinity assembly statistics. From the assembly, the total number of transcripts generated were 34170 with a minimum contig length of 101 bp and a maximum of 9227 bp. The average contig length was 673 bp with GC content of 62.47 %.

**Figure 1.**
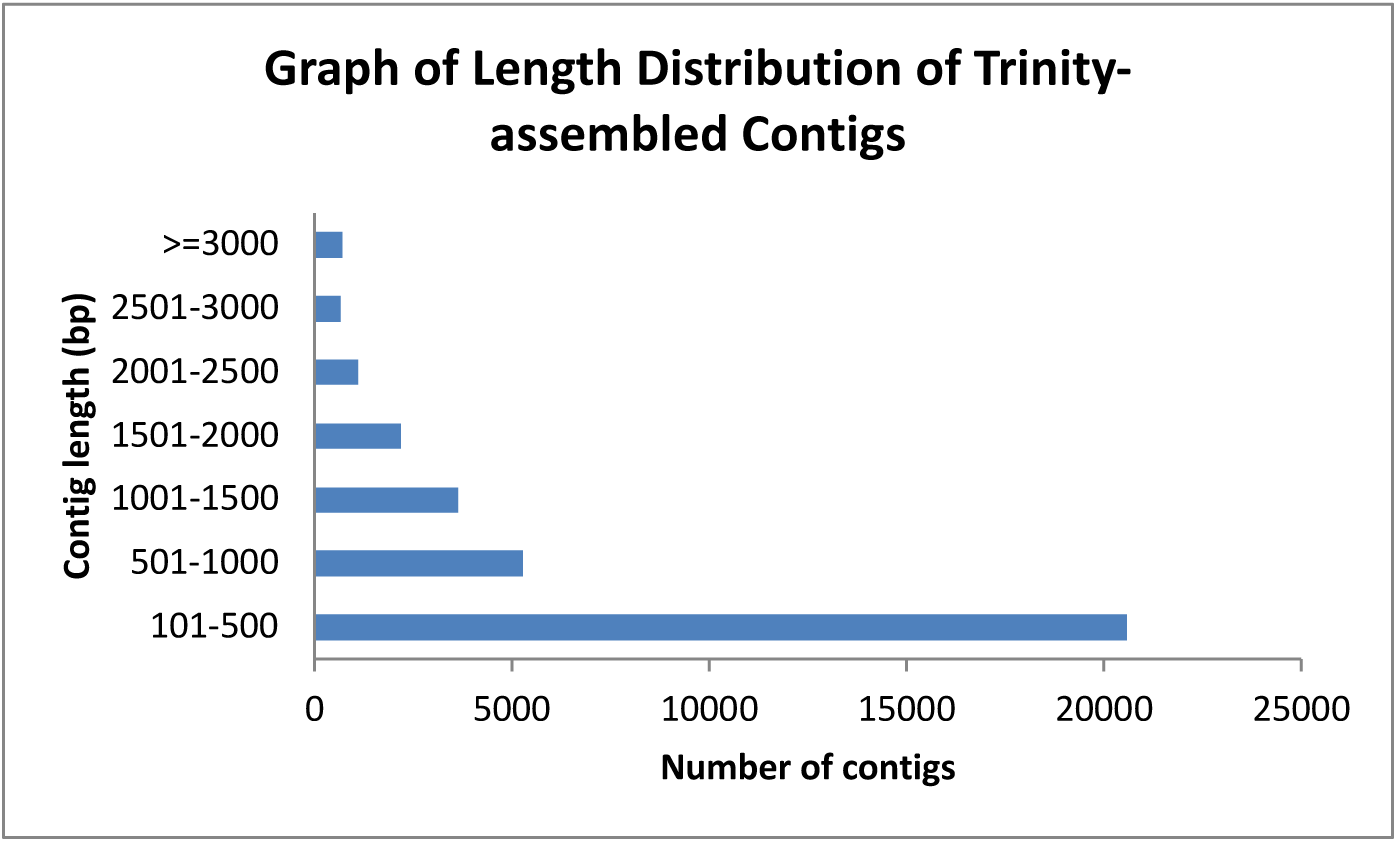
Graph showing the length distribution of Trinity-assembled contigs. A total of 34170 contigs were assembled. The most abundant contig lengths in the assembly were in the range of 101 bp to 500 bp while the least abundant were in the range of 2501 bp to 3000 bp.

### Differentially expressed transcripts and functional annotation

Three pairwise comparisons of the transcriptomes were done, which were 15 °C vs 5 °C, 5 °C vs 0 °C and 15 °C vs 21 °C. A total of 133 transcripts were identified to be at least two-fold differentially expressed, with several overlaps among the three pairwise comparisons (Table 3-5). The threshold was set with false discovery-corrected statistical significance at most 0.001 based on its normalized expression value measured in FPKM (fragments per feature kilobase per million reads mapped). However, the RNA-Seq count data were normalized using the TMM normalization method before the FPKM values were generated. The differentially expressed transcripts are described in Table 3-5.

**Table 3.** Brief description based on functional groups of 45 transcripts represented in Figure 2(a-c), where *Rhodotorula* sp. USM-PSY62 was exposed to temperature shift from 15 °C to 5 °C.

| No. | Transcript ID | Log fold change | Expression | eggNOG predictions (Accession\|Name\|[Category]) |
| --- | --- | --- | --- | --- |
| ***Cellular processes and signalling*** |  |  |  |  |
| 1 | comp2729_c0_seq1 | -2.643854 | down | COG0330 \|HflC, Membrane protease subunits, stomatin/prohibitin homologs [Posttranslational modification] |
| 2 | comp2184_c0_seq1 | -1.263762 | down | COG0702 \|COG0702, Predicted nucleoside-diphosphate-sugar epimerases[Cell wall/membrane/envelope/biogenesis] |
| 3 | comp6513_c0_seq1 | -1.493231 | down | COG0071 \|IbpA, Molecular chaperone (small heat shock protein)[Posttranslational modification] |
| 4 | comp15275_c0_seq1 | -1.695049 | down | COG5074 \|COG5074, t-SNARE complex subunit, syntaxin[Intracellular trafficking,secretion and vesicular transport] |
| 5 | comp12847_c0_seq1 | 9.705356 | up | COG4238 \|COG4238, Murein lipoprotein [Cell wall/membrane/envelope/biogenesis] |
| 6 | comp6405_c0_seq1 | -6.931226 | down | COG5052 \|YOP1, Protein involved in membrane traffic [Intracellular trafficking, secretion and vesicular transport] |
| 7 | comp8473_c0_seq9 | -2.430342 | down | COG0642 \|BaeS, Signal transduction histidine kinase [Signal transduction mechanism] |
| ***Information storage and processing*** |  |  |  |  |
| 8 | comp7487_c0_seq2 | 8.018526 | up | COG1670 \|RimL, Acetyltransferases, including N-acetylases of ribosomal proteins [Translation, ribosomal structure and biogenesis] |
| 9 | comp8916_c0_seq6 | -6.37717 | down | COG0507 \|RecD, ATP-dependent exoDNase (exonuclease V), alpha subunit - helicase superfamily I member [Replication, recombination] |
|  |  |  |  | and repair] |
| 10 | comp8916_c0_seq10 | 1.746675 | up | COG0507 \|RecD, ATP-dependent<br>exoDNase (exonuclease V), alpha<br>subunit - helicase superfamily I<br>member [Replication, recombination<br>and repair] |
| 11 | comp8916_c0_seq3 | 3.015206 | up | COG0507 \|RecD, ATP-dependent<br>exoDNase (exonuclease V), alpha<br>subunit - helicase superfamily I<br>member [Replication, recombination<br>and repair] |
| 12 | comp8916_c0_seq1 | -8.012335 | down | COG0507 \|RecD, ATP-dependent<br>exoDNase (exonuclease V), alpha<br>subunit - helicase superfamily I<br>member [Replication, recombination<br>and repair] |
| 13 | comp8936_c0_seq12 | 9.042724 | up | COG0553 \|HepA, Superfamily II<br>DNA/RNA helicases, SNF2 family<br>[Transcription] |
| 14 | comp8936_c0_seq4 | 2.409688 | up | COG0553 \|HepA, Superfamily II<br>DNA/RNA helicases, SNF2 family<br>[Transcription] |
| 15 | comp8191_c0_seq3 | 2.876829 | up | COG0553 \|HepA, Superfamily II<br>DNA/RNA helicases, SNF2 family<br>[Transcription] |
| 16 | comp8936_c0_seq8 | -1.634288 | down | COG0553 \|HepA, Superfamily II<br>DNA/RNA helicases, SNF2 family<br>[Transcription] |
| 17 | comp8739_c1_seq3 | 1.688598 | up | COG5179 \|TAF1, Transcription<br>initiation factor TFIID, subunit<br>TAF1 [Transcription] |
| 18 | comp5490_c0_seq1 | -1.228738 | down | COG5084 \|YTH1, Cleavage and<br>polyadenylation specificity factor<br>(CPSF) Clipper subunit and related<br>makorin family Zn- finger proteins<br>[Transcription] |
| 19 | comp2135_c0_seq2 | 1.620837 | up | COG5147 \|REB1, Myb superfamily<br>proteins, including transcription<br>factors and mRNA splicing factors<br>[Transcription] |
| 20 | comp5468_c0_seq1 | -5.856946 | down | COG5169 \|HSF1, Heat shock<br>transcription factor [Transcription] |
| 21 | comp8426_c1_seq6 | 2.20702 | up | COG5256 TEF1, Translation elongation factor EF-1alpha (GTPase) [Translation, ribosomal structure and biogenesis] |
| **Metabolism** |  |  |  |  |
| 22 | comp6898_c1_seq2 | -1.841753 | down | COG0477 \|ProP, Permeases of the major facilitator superfamily [Carbohydrate transport and metabolism] |
| 23 | comp4572_c0_seq1 | -2.040901 | down | COG0477 \|ProP, Permeases of the major facilitator superfamily [Carbohydrate transport and metabolism] |
| 24 | comp11718_c0_seq1 | -1.4509 | down | COG1028 \|FabG, Dehydrogenases with different specificities (related to short-chain alcohol dehydrogenases) [Secondary metabolites biosynthesis, transport and catabolism] |
| 25 | comp6796_c0_seq1 | -1.300862 | down | COG1902 \|NemA, NADH:flavin oxidoreductases, Old Yellow Enzyme family [Energy production and conversion] |
| 26 | comp7209_c0_seq2 | -2.670481 | down | COG2273 \|SKN1, Beta-glucanase/Beta-glucan synthetase [Carbohydrate transport and metabolism] |
| 27 | comp7230_c0_seq2 | -8.091322 | down | COG1893 \|ApbA, Ketopantoate reductase [Coenzyme transport and metabolism] |
| 28 | comp13825_c0_seq1 | -1.331793 | down | COG1028 \|FabG, Dehydrogenases with different specificities (related to short-chain alcohol dehydrogenases) [Secondary metabolites biosynthesis, transport and catabolism] |
| 29 | comp8089_c0_seq4 | -1.615364 | down | COG0604 \|Qor, NADPH:quinone reductase and related Zn-dependent oxidoreductases [Energy production and conversion] |
| 30 | comp7629_c0_seq3 | -5.659021 | down | COG0604 \|Qor, NADPH:quinone reductase and related Zn-dependent oxidoreductases [Energy production and conversion] |
| 31 | comp14257_c0_seq1 | -1.210936 | down | COG0807 RibA, GTP cyclohydrolase II [Coenzyme transport and metabolism] |
| 32 | comp8113_c0_seq6 | 1.471048 | up | COG1785 \|PhoA, Alkaline phosphatase [Inorganic ion transport and metabolism] |
| 33 | comp6337_c0_seq1 | -2.091992 | down | COG2124 \|CypX, Cytochrome P450 [Secondary metabolites biosynthesis, transport and catabolism] |
| 34 | comp3495_c0_seq2 | -1.842732 | down | COG3250 \|LacZ, Beta-galactosidase/beta-glucuronidase [Carbohydrate transport and metabolism] |
| 35 | comp8089_c0_seq3 | -2.837434 | down | COG0604 \|Qor, NADPH:quinone reductase and related Zn-dependent oxidoreductases [Energy production and conversion] |
| 36 | comp3523_c0_seq1 | 1.597053 | up | COG0534 \|NorM, Na<sup>+</sup>-driven multidrug efflux pump [Inorganic ion transport and metabolism] |
| 37 | comp8278_c0_seq4 | 6.999321 | up | COG0598 \|CorA, Mg<sup>2+</sup> and Co<sup>2+</sup> transporters [Inorganic ion transport and metabolism] |
| 38 | comp7430_c0_seq2 | -3.390306 | down | COG0604 \|Qor, NADPH:quinone reductase and related Zn-dependent oxidoreductases [Energy production and conversion] |
| 39 | comp8366_c0_seq2 | -4.041479 | down | COG1472 \|BglX, Beta-glucosidase-related glycosidases [Carbohydrate transport and metabolism] |
| 40 | comp5954_c0_seq2 | -2.249075 | down | COG0408 \|HemF, Coproporphyrinogen III oxidase [Coenzyme transport and metabolism] |
| 41 | comp8366_c0_seq4 | -3.486855 | down | COG1472 \|BglX, Beta-glucosidase-related glycosidases [Carbohydrate transport and metabolism] |
| 42 | comp6452_c0_seq1 | -1.474342 | down | COG0807 \|RibA, GTP cyclohydrolase II [Coenzyme transport and metabolism] |
| 43 | comp6069_c1_seq1 | -1.380557 | down | COG2072 \|TrkA, Predicted flavoprotein involved in K<sup>+</sup> transport [Inorganic ion transport and metabolism] |
| 44 | comp8844_c0_seq2 | 2.812792 | up | COG2937 \|PlsB, Glycerol-3- |
|  |  |  |  | phosphate O-acyltransferase [Lipid transport and metabolism] |
| 45 | comp2593_c0 | 2.6337254 | up | NOG27164 \|Catalase [CAT, EC 1.11.1.6] |

**Table 4.** Brief description based on functional groups of 47 transcripts represented in Figure 3(a-c), where *Rhodotorula* sp. USM-PSY62 was exposed to temperature shift from 5 °C to 0 °C.

| No. | Transcript ID | Log fold change | Expression | eggNOG predictions (Accession\|Name\|[Category]) |
| --- | --- | --- | --- | --- |
| ***Cellular processes and signalling*** |  |  |  |  |
| 1 | comp6513_c0_seq1 | 1.129581 | up | COG0071 \|IbpA, Molecular chaperone (small heat shock protein) [Posttranslational modification] |
| 2 | comp3254_c1_seq1 | -1.018463 | down | COG0326 \|HtpG, Molecular chaperone, HSP90 family [Posttranslational modification] |
| 3 | comp6918_c0_seq1 | -8.803772 | down | COG1196 \|Smc, Chromosome segregation ATPases [Cell cycle control, cell division, chromosome partitioning] |
| 4 | comp6405_c0_seq1 | -1.387866 | down | COG5052 \|YOP1, Protein involved in membrane traffic [Intracellular trafficking, secretion and vesicular transport] |
| 5 | comp4593_c0_seq2 | 1.176855 | up | COG1215 \|COG1215, Glycosyltransferases, probably involved in cell wall biogenesis [Cell wall/membrane/envelope/biogenesis] |
| 6 | comp7922_c4_seq2 | 7.138811 | up | COG5391 \|COG5391, Phox homology (PX) domain protein [Intracellular trafficking, secretion and vesicular transport] |
| 7 | comp8473_c0_seq9 | 1.93212 | up | COG0642 \|BaeS, Signal transduction histidine kinase [Signal transduction mechanism] |
| 8 | comp7948_c0_seq3 | 1.251941 | up | COG0642 \|BaeS, Signal transduction histidine kinase [Signal transduction mechanism] |
| ***Information storage and processing*** |  |  |  |  |
| 9 | comp7487_c0_seq2 | -5.560906 | down | COG1670 \|RimL, Acetyltransferases, including N-acetylases of ribosomal proteins [Translation, ribosomal structure] |
|  |  |  |  | and biogenesis] |
| 10 | comp8916_c0_seq6 | -1.868045 | down | COG0507 \|RecD, ATP-dependent<br>exoDNase (exonuclease V), alpha<br>subunit - helicase superfamily I<br>member [Replication,<br>recombination and repair] |
| 11 | comp8916_c0_seq10 | 6.261352 | up | COG0507 \|RecD, ATP-dependent<br>exoDNase (exonuclease V), alpha<br>subunit – helicase superfamily I<br>member [Replication,<br>recombination and repair] |
| 12 | comp8916_c0_seq1 | 1.590589 | up | COG0507 \|RecD, ATP-dependent<br>exoDNase (exonuclease V), alpha<br>subunit - helicase superfamily I<br>member [Replication,<br>recombination and repair] |
| 13 | comp8936_c0_seq12 | -7.436077 | down | COG0553\|HepA, Superfamily II<br>DNA/RNA helicases, SNF2 family<br>[Transcription] |
| 14 | comp7744_c0_seq3 | 7.384942 | up | COG0553 \|HepA, Superfamily II<br>DNA/RNA helicases, SNF2 family<br>[Transcription] |
| 15 | comp6218_c0_seq1 | -2.007918 | down | COG5169 \|HSF1, Heat shock<br>transcription factor [Transcription] |
| 16 | comp9006_c1_seq2 | 2.325696 | up | COG5211 \|SSU72, RNA<br>polymerase II-interacting protein<br>involved in transcription start site<br>selection [Transcription] |
| 17 | comp8936_c0_seq4 | -2.195524 | down | COG0553 \|HepA, Superfamily II<br>DNA/RNA helicases, SNF2 family<br>[Transcription] |
| 18 | comp8191_c0_seq3 | 6.03144 | up | COG0553 \|HepA, Superfamily II<br>DNA/RNA helicases, SNF2 family<br>[Transcription] |
| 19 | comp8936_c0_seq8 | 10.16288 | up | COG0553 \|HepA, Superfamily II<br>DNA/RNA helicases, SNF2 family<br>[Transcription] |
| 20 | comp8739_c1_seq3 | 5.647731 | up | COG5179 \|TAF1, Transcription<br>initiation factor TFIID, subunit<br>TAF1 [Transcription] |
| 21 | comp5490_c0_seq1 | -1.547657 | down | COG5084 YTH1, Cleavage and<br>polyadenylation specificity factor<br>(CPSF) Clipper subunit and related makorin family Zn-finger proteins<br>[Transcription] |
| 22 | comp5468_c0_seq1 | -2.241933 | down | COG5169 \|HSF1, Heat shock<br>transcription factor [Transcription] |
| ***Metabolism*** |  |  |  |  |
| 23 | comp2909_c1_seq1 | 3.184432 | up | COG0477 \|ProP, Permeases of the<br>major facilitator superfamily<br>[Carbohydrate transport and<br>metabolism] |
| 24 | comp8176_c0_seq2 | 1.43372 | up | COG0477 \|ProP, Permeases of the<br>major facilitator superfamily<br>[Carbohydrate transport and<br>metabolism] |
| 25 | comp6898_c1_seq2 | 1.137336 | up | COG0477 \|ProP, Permeases of the<br>major facilitator superfamily<br>[Carbohydrate transport and<br>metabolism] |
| 26 | comp4572_c0_seq1 | 1.063855 | up | COG0477 \|ProP, Permeases of the<br>major facilitator superfamily<br>[Carbohydrate transport and<br>metabolism] |
| 27 | comp8732_c0_seq1 | 1.720333 | up | COG0477 \|ProP, Permeases of the<br>major facilitator superfamily<br>[Carbohydrate transport and<br>metabolism] |
| 28 | comp11718_c0_seq1 | 1.589077 | up | COG1028 \|FabG, Dehydrogenases<br>with different specificities (related<br>to short-chain alcohol<br>dehydrogenases) [Secondary<br>metabolites biosynthesis, transport<br>and catabolism] |
| 29 | comp7209_c0_seq2 | 3.95132 | up | COG2273 \|SKN1, Beta-glucanase/Beta-glucan synthetase<br>[Carbohydrate transport and<br>metabolism] |
| 30 | comp7230_c0_seq2 | -1.822073 | down | COG1893 \|ApbA, Ketopantoate<br>reductase [Coenzyme transport and<br>metabolism] |
| 31 | comp13825_c0_seq1 | 1.839644 | up | COG1028 FabG, Dehydrogenases<br>with different specificities (related<br>to short-chain alcohol<br>dehydrogenases) [Secondary<br>metabolites biosynthesis, transport and catabolism] |
| 32 | comp8089_c0_seq4 | 2.621868 | up | COG0604 \|Qor, NADPH:quinone reductase and related Zn-dependent oxidoreductases [Energy production and conversion] |
| 33 | comp14257_c0_seq1 | -1.192267 | down | COG0807 \|RibA, GTP cyclohydrolase II [Coenzyme transport and metabolism] |
| 34 | comp8113_c0_seq6 | -2.119545 | down | COG1785 \|PhoA, Alkaline phosphatase [Inorganic ion transport and metabolism] |
| 35 | comp9040_c0_seq32 | 7.775199 | up | COG2303 \|BetA, Choline dehydrogenase and related flavoproteins [Amino acid transport and metabolism] |
| 36 | comp3495_c0_seq2 | 2.738382 | up | COG3250 \|LacZ, Beta-galactosidase/beta-glucuronidase [Carbohydrate transport and metabolism] |
| 37 | comp8089_c0_seq3 | 2.604183 | up | COG0604 \|Qor, NADPH:quinone reductase and related Zn-dependent oxidoreductases [Energy production and conversion] |
| 38 | comp3523_c0_seq1 | 6.589149 | up | COG0534 \|NorM, Na<sup>+</sup>-driven multidrug efflux pump [Inorganic ion transport and metabolism] |
| 39 | comp8278_c0_seq4 | -8.863971 | down | COG0598 \|CorA, Mg<sup>2+</sup> and Co<sup>2+</sup> transporters [Inorganic ion transport and metabolism] |
| 40 | comp7430_c0_seq2 | 1.253994 | up | COG0604 \|Qor, NADPH:quinone reductase and related Zn-dependent oxidoreductases [Energy production and conversion] |
| 41 | comp8089_c0_seq2 | 3.820099 | up | COG0604 \|Qor, NADPH:quinone reductase and related Zn-dependent oxidoreductases [Energy production and conversion] |
| 42 | comp8366_c0_seq2 | 8.047347 | up | COG1472 \|BglX, Beta-glucosidase-related glycosidases [Carbohydrate transport and metabolism] |
| 43 | comp8592_c0_seq12 | 5.558765 | up | COG0531 \|PotE, Amino acid transporters [Amino acid transport and metabolism] |
| 44 | comp8366_c0_seq4 | 5.263325 | up | COG1472 \|BglX, Beta-glucosidase-related glycosidases [Carbohydrate transport and metabolism] |
| 45 | comp6452_c0_seq1 | 1.099371 | up | COG0807 \|RibA, GTP cyclohydrolase II [Coenzyme transport and metabolism] |
| 46 | comp6069_c1_seq1 | -1.044898 | down | COG2072 \|TrkA, Predicted flavoprotein involved in K<sup>+</sup> transport [Inorganic ion transport and metabolism] |
| 47 | comp2593_c0 | -2.956503 | down | NOG27164 \|Catalase [CAT, EC 1.11.1.6] |

**Table 5.**
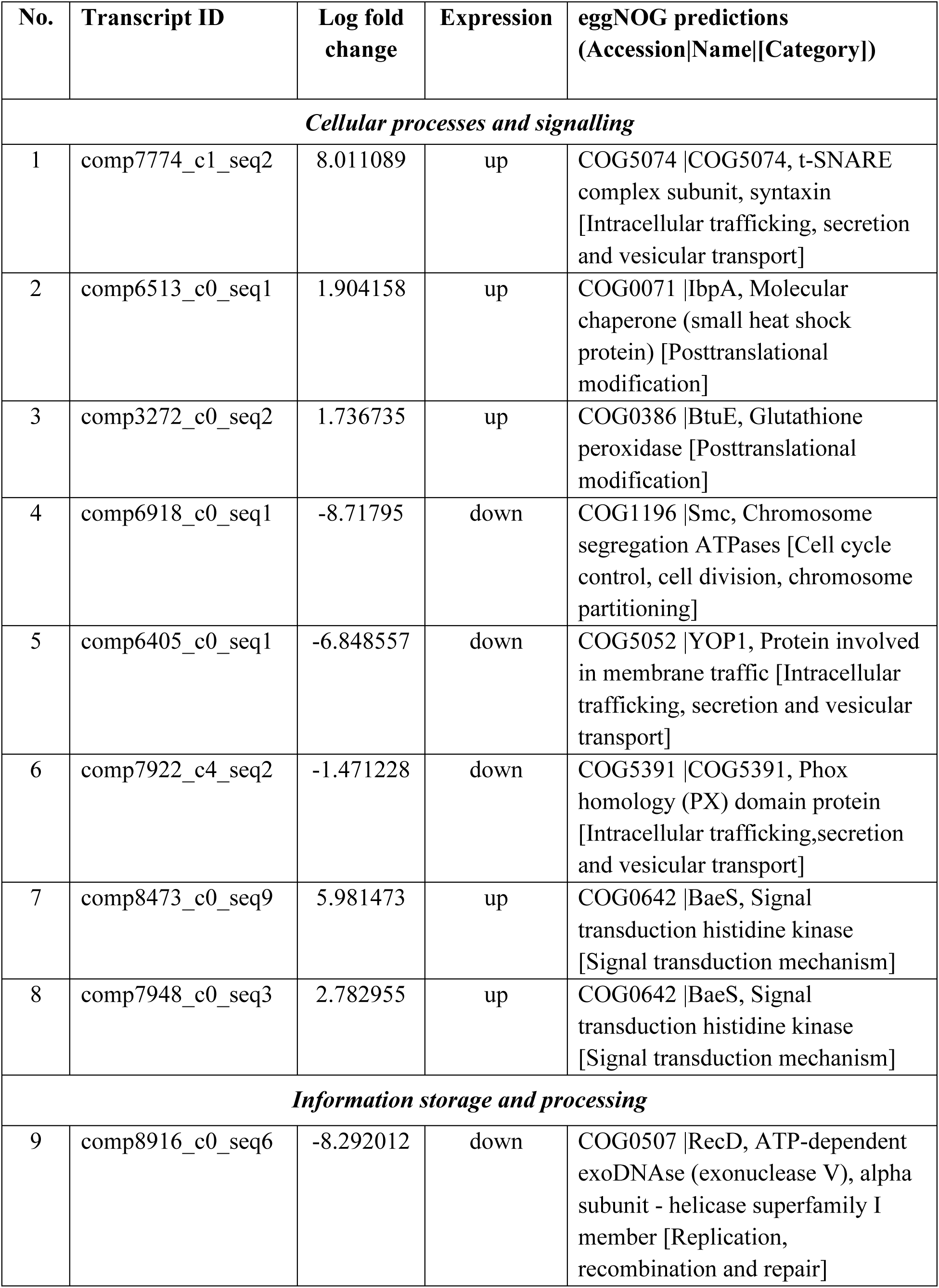
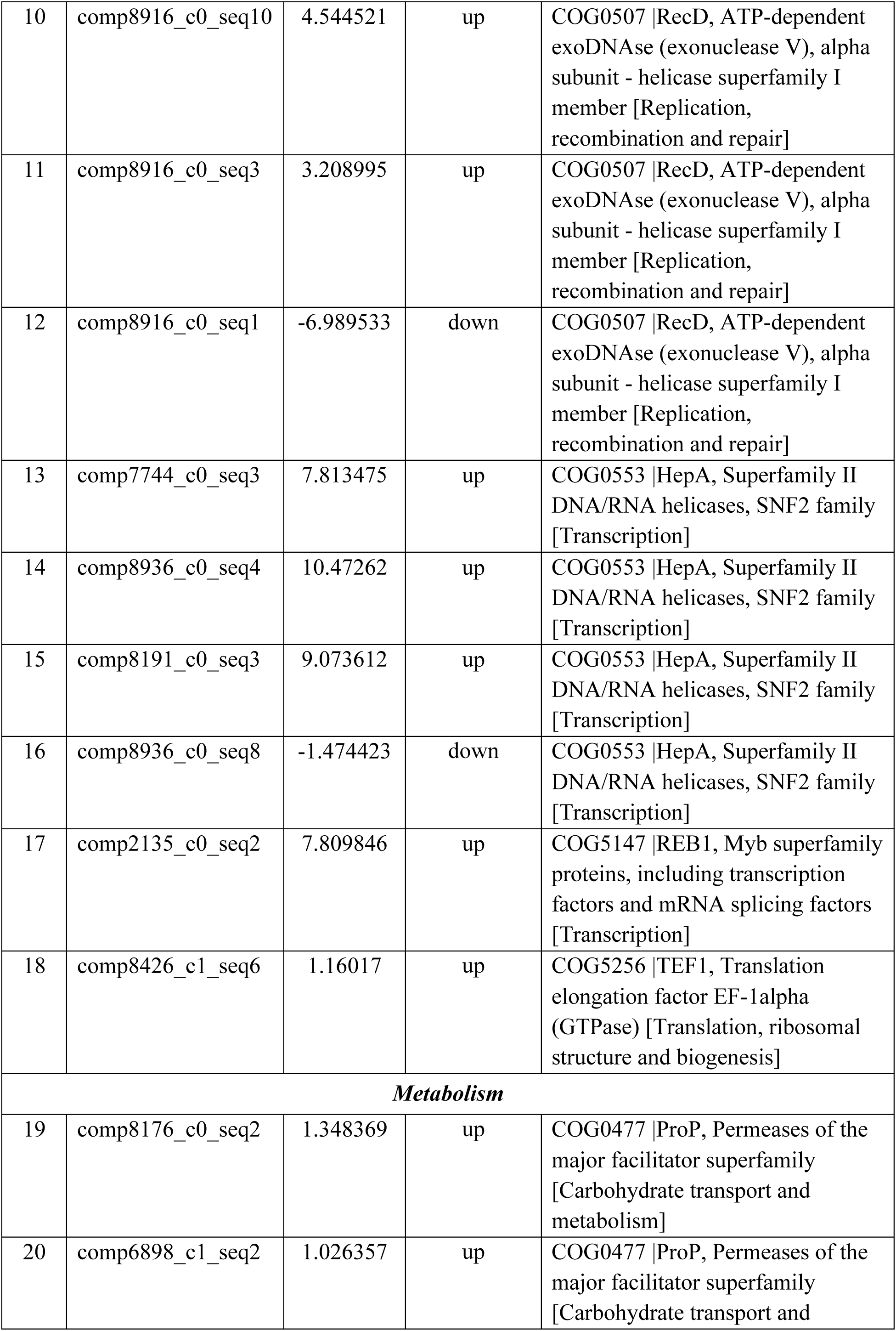
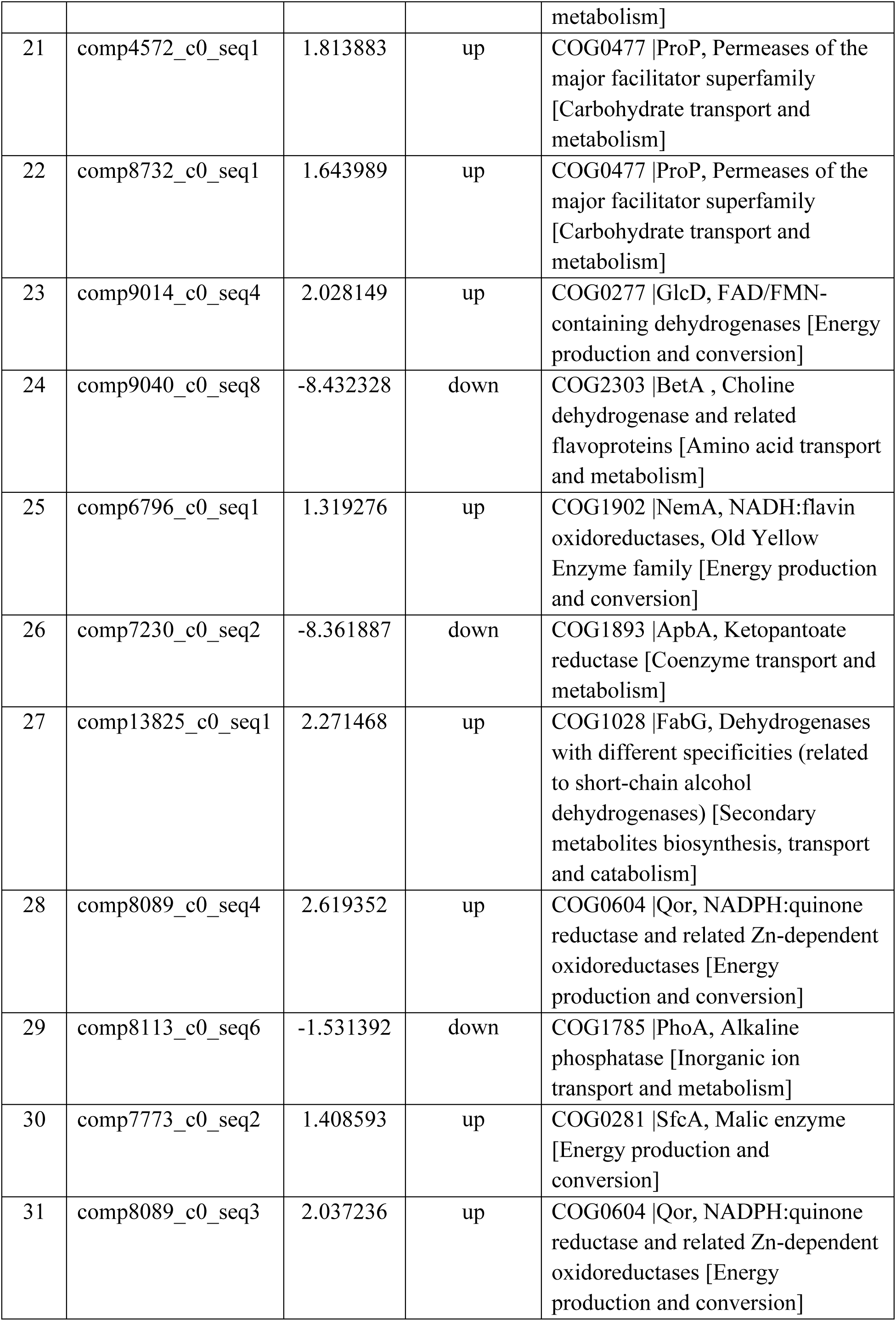
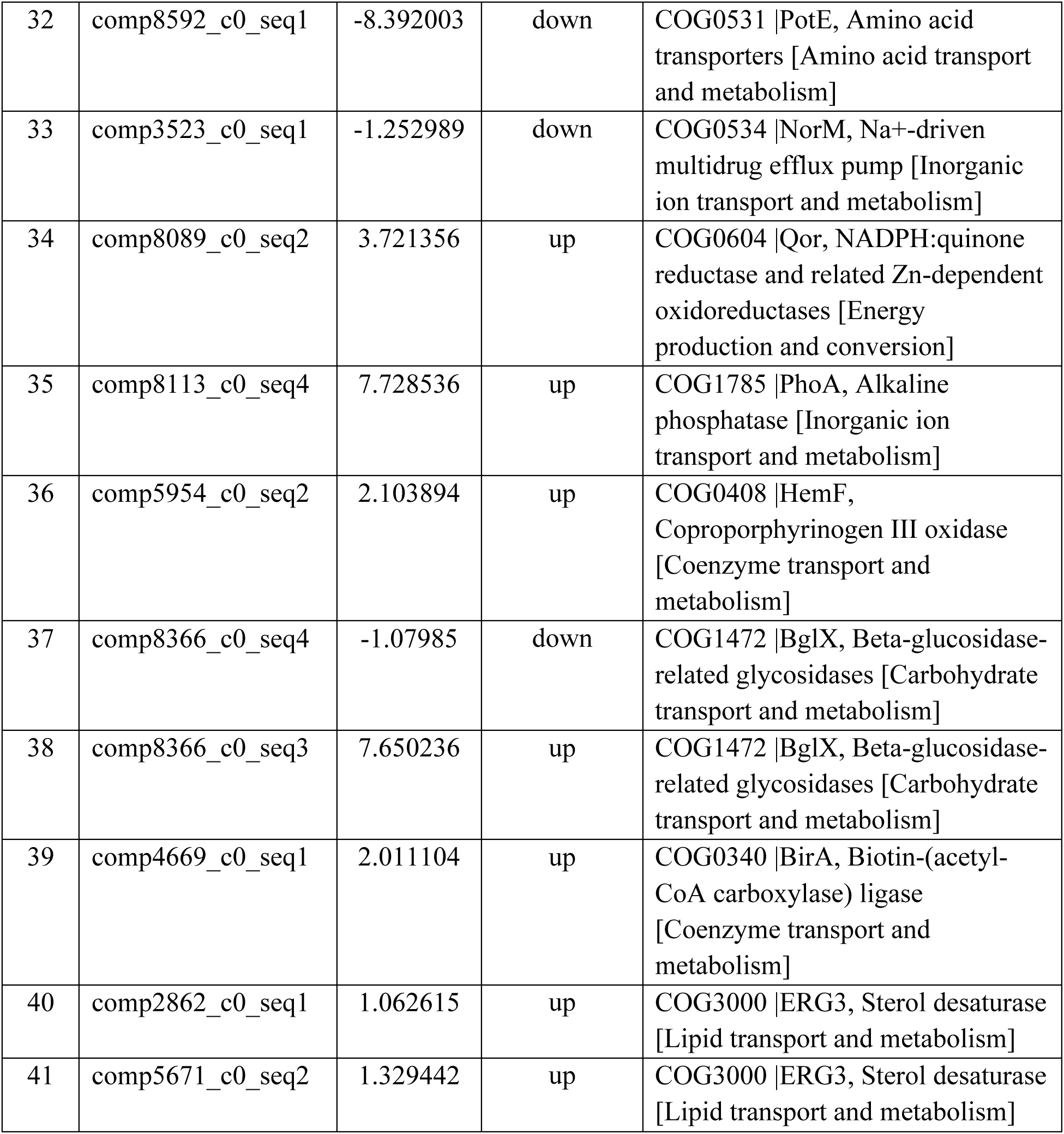
List of 41 transcripts that were differentially expressed by at least two-fold under condition from 15 °C to 21 °C, sorted according to eggNOG predictions.

The qRT-PCR validation graph is given in Online Resource 1. Based on the transcriptomic analysis, five transcripts with significant up- or down-regulation in expression were chosen for qRT-PCR validation. The relative expression of each transcript was normalized against the expression of a control gene, actin, and analyzed using Livak’s (2^-ΔΔCq^) method. To validate the transcriptomic analysis, the qRT-PCR result was compared against it. As an example, the down-regulated expression of HSF1 (comp5468_c0_seq1; Table 3; Figure 2b) by a log2 fold change value of −5.86 (15 °C to 5 °C) was similarly illustrated by qRT-PCR technique, which showed a log2 fold change value of approximately −3.00 (Online Resource 1).

**Figure 2.**
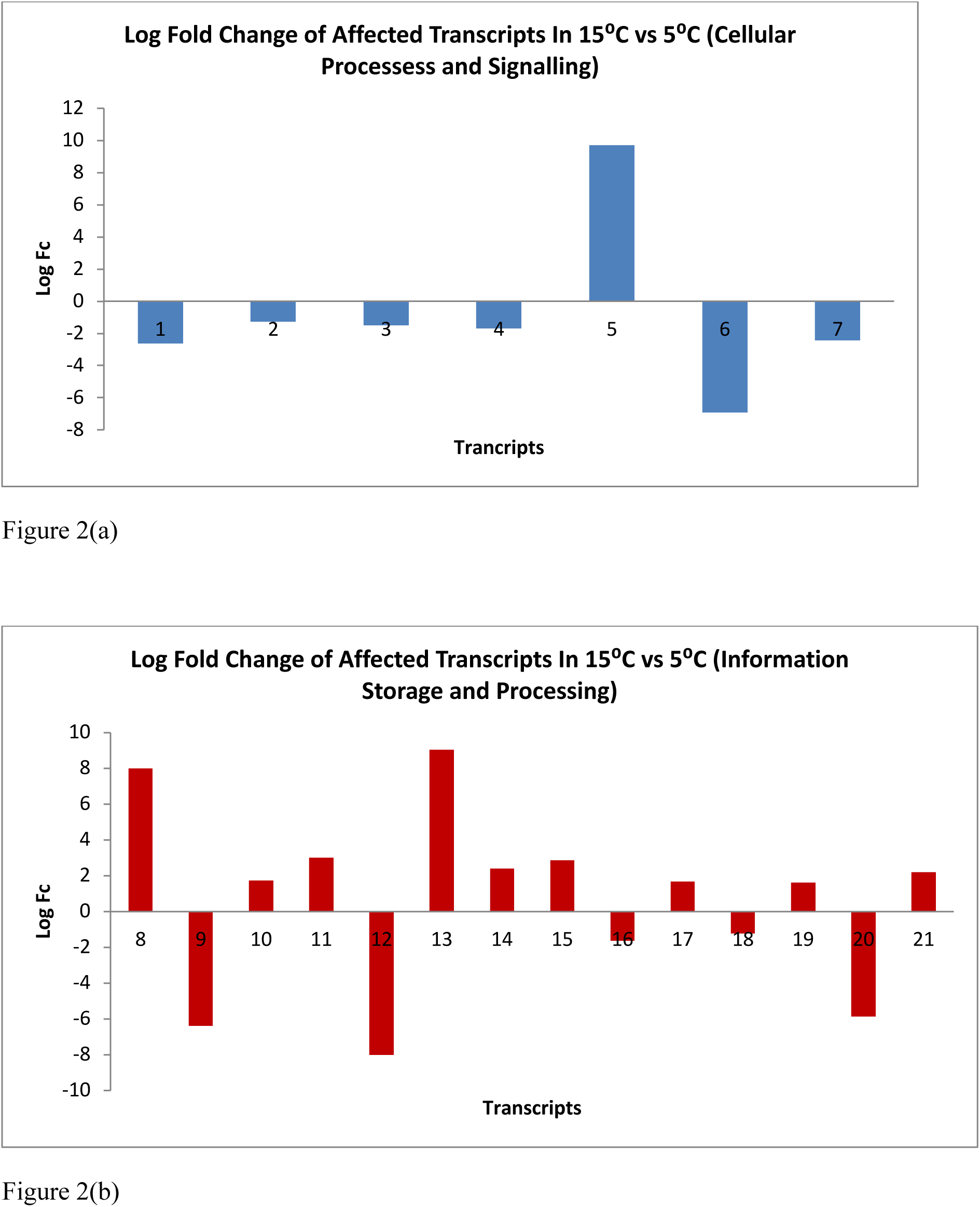
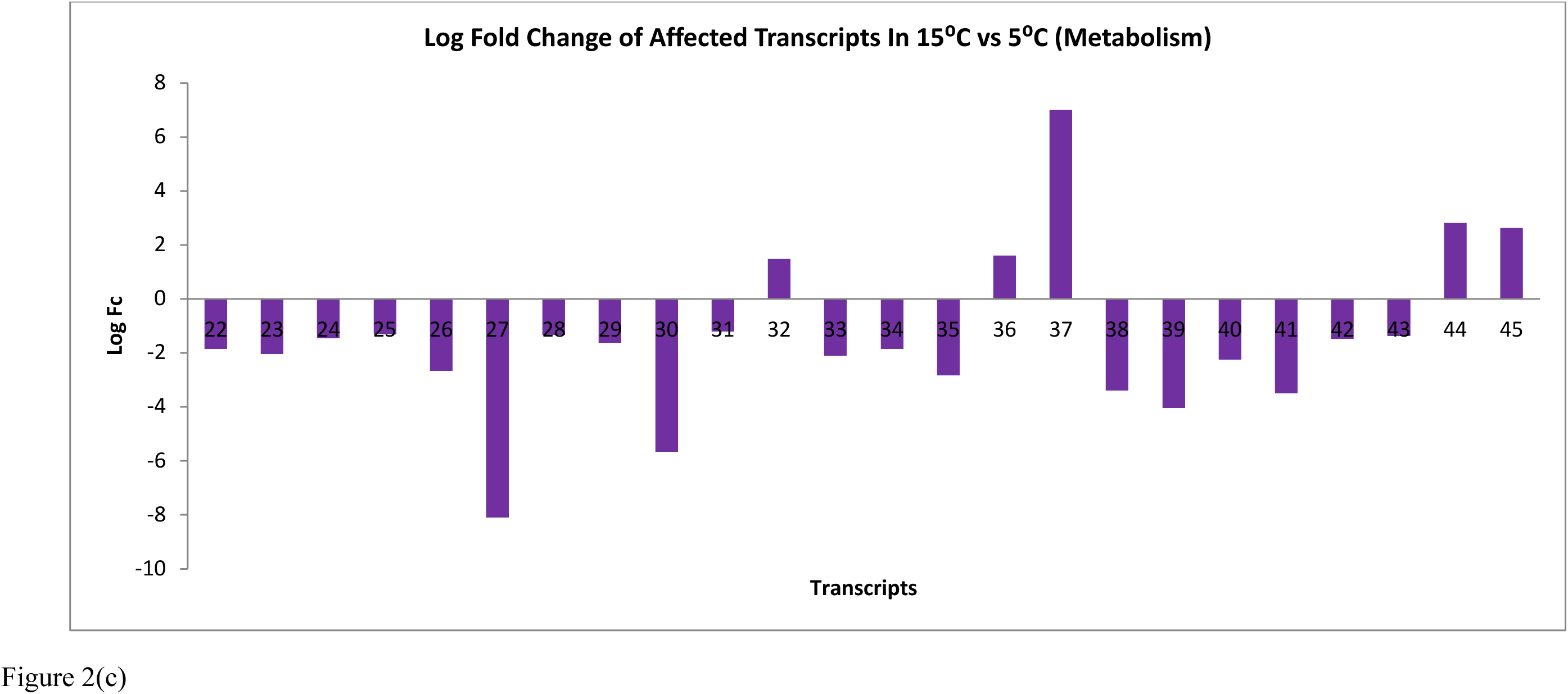
Graphs showing log fold change of transcripts expression when temperature shifted from 15 °C to 5 °C. Figure 2(a) shows affected transcripts under the “Cellular Processes and Signalling” category, 2(b) shows affected transcripts under “Information Storage and Processing” and 2(c) shows affected transcripts under “Metabolism”. Description on each transcript is listed in Table 3.

The differentially expressed transcripts were classified into three main groups based on the eggNOG annotations, namely ‘cellular processing and signaling’, ‘information storage and processing’, and finally ‘metabolism’.

Bar graphs representing detailed response of every differentially expressed transcripts between cold adaptation responses (15 °C to 5 °C and 5 °C to 0 °C) and heat adaptation responses (15 °C to 21 °C) are shown in Figure 2, Figure 3 and Figure 4, respectively.

**Figure 3.**
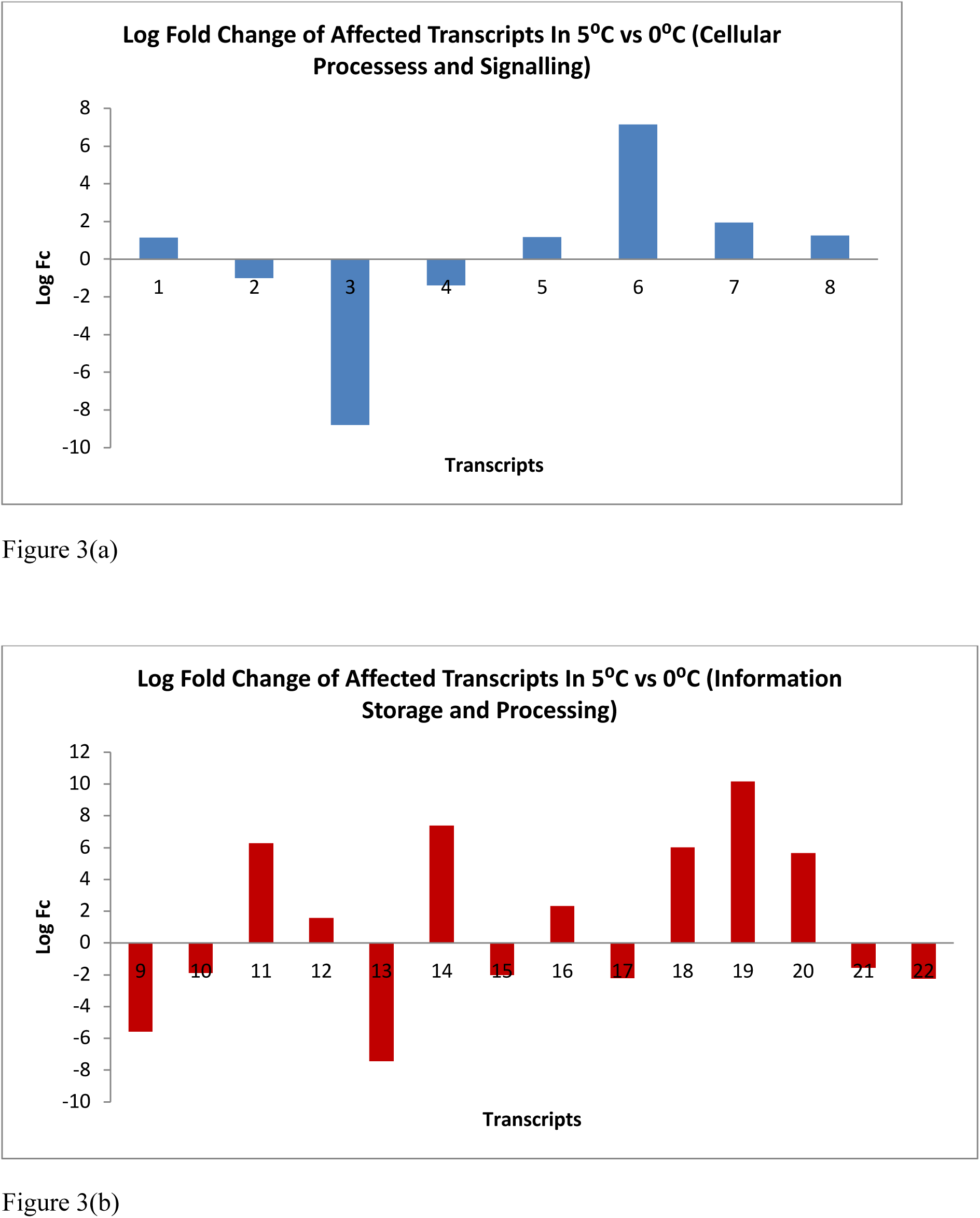
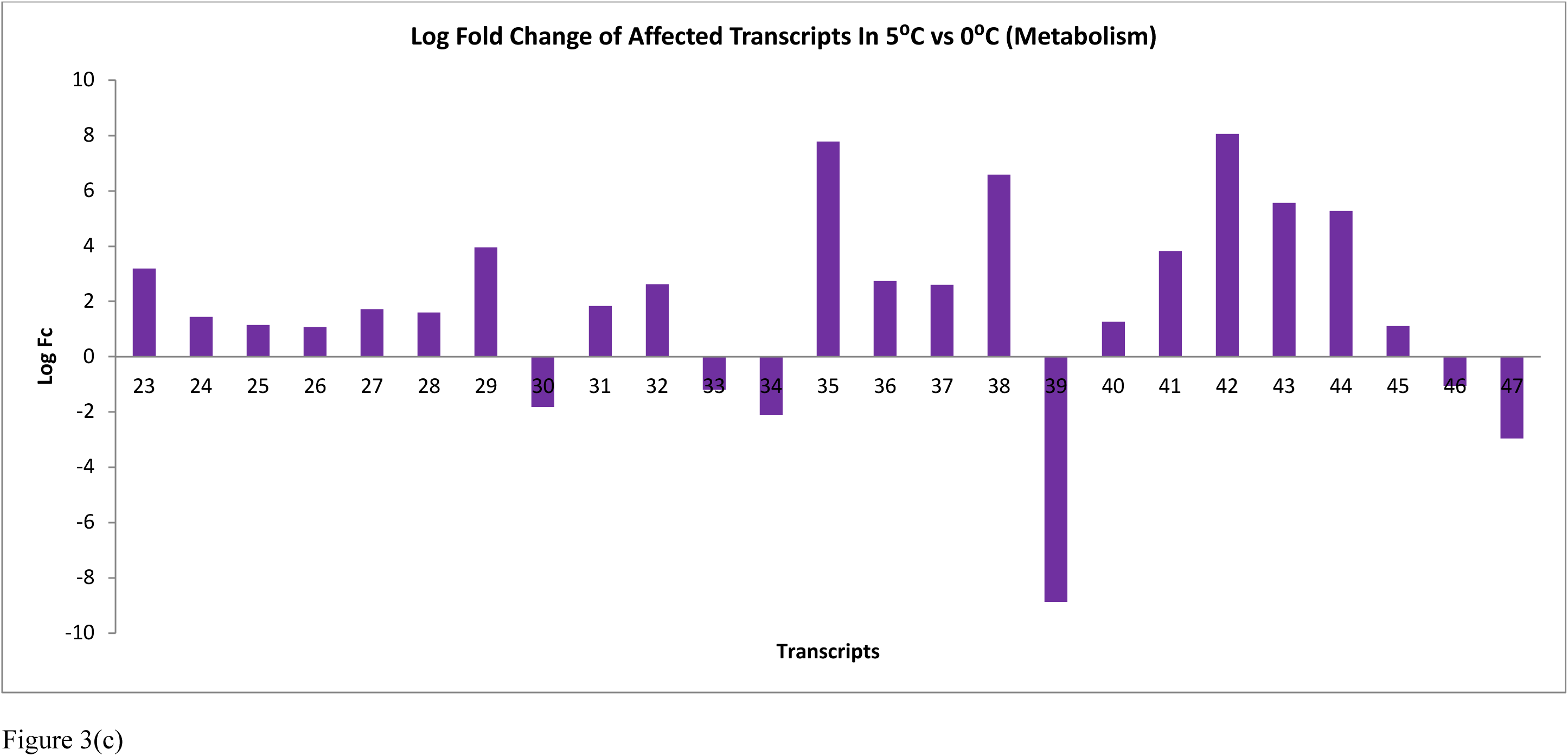
Graphs showing log fold change of transcripts expression when temperature shifted from 5 °C to 0 °C. Figure 3(a) shows affected transcripts under the “Cellular Processes and Signalling” category, 3(b) shows affected transcripts under “Information Storage and Processing” and 3(c) shows affected transcripts under “Metabolism”. Description on each transcript is listed in Table 4.

**Figure 4.**
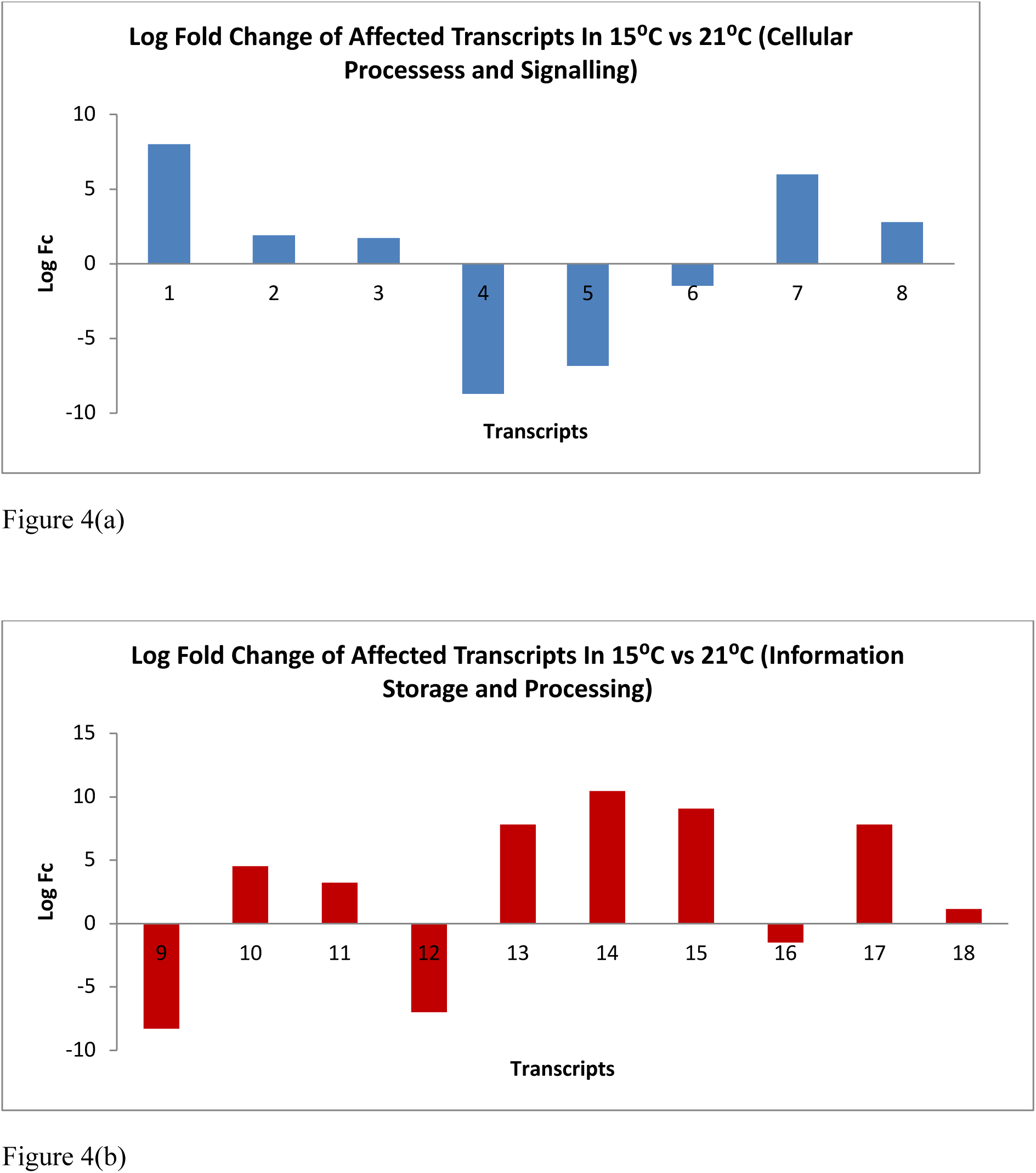
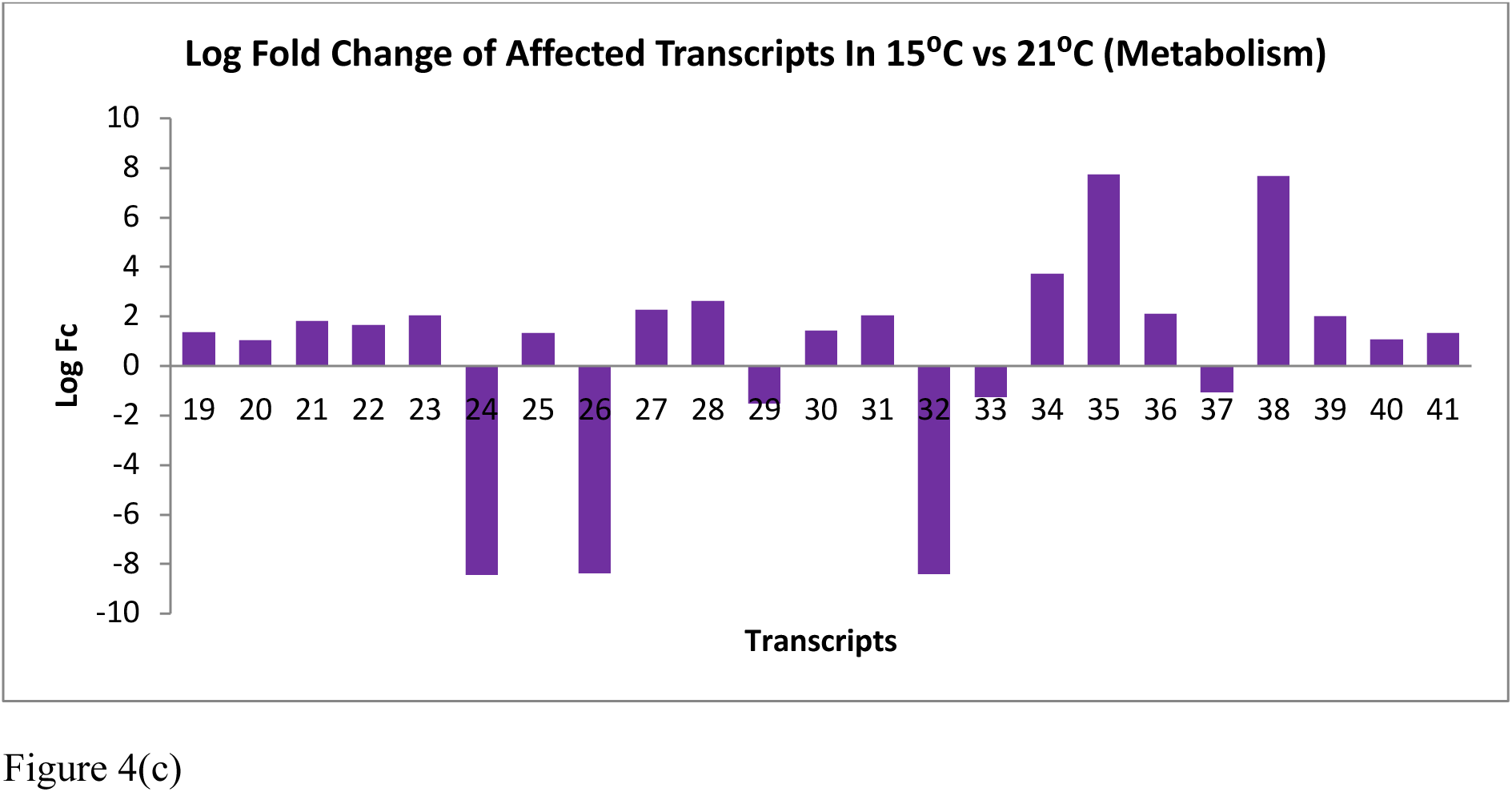
Graphs showing log fold change of transcripts expression when temperature shifted from 15 °C to 21 °C. Figure 4(a) shows affected transcripts under the “Cellular Processes and Signalling” category, 4(b) shows affected transcripts under “Information Storage and Processing” and 4(c) shows affected transcripts under “Metabolism”. Description on each transcript is listed in Table 5.

**Table 6** Brief description based on functional groups of 41 transcripts represented in Figure 4(a-c), where *Rhodotorula* sp. USM-PSY62 was exposed to temperature shift from 15 °C to 21 °C.

## Discussion

In order to survive, organisms must be able to acclimatize when confronted with stressful environmental circumstances. A common form of stress in an ecosystem is changes in its ambient temperature which interferes with the state of metabolism of its inhabitants. These populations of inhabitants will need to adapt in order to flourish in the new settings (Hoffman 2000). The effects produced are mostly on the chemical and enzyme-based reactions inside the organisms as well as structural changes of their cellular constituents. Temperature shifts also elicit complex cellular responses involving genes. Transcriptional reactions engage many genes that can be categorized according to the external variations that were imposed.

The impact of environmental change on organisms is strongly influenced by their ability to adaptably adjust their physiology. As a psychrophilic yeast, *Rhodotorula* sp. USM-PSY62 should similarly be able to regulate its response to varying environmental conditions. The physiological perturbations and pressures inflicted upon it should reveal distinctive adaptive strategies necessary for its survival. In this study, the cold-shock adaptive strategy was triggered by shifting the temperature from the optimum of 15 °C to 5 °C and 0 °C. The heat-shock response was also investigated by transferring the temperature from 15 °C to 21 °C. Psychrophilic microorganisms up- or down-regulate the expression of a number of genes in response to sudden change in growth temperatures (De Maayer et al. 2014). Such physiological responses normally begin with recognition of the temperature stress. The perceived stress signals are then transmitted to the cellular machinery to activate the adaptive responses (Xiong et al. 2002).

The *YOP1* gene of USM-PSY62 was the most down-regulated in expression under cold shock conditions with log2 fold change values of −6.93 (comp6405_c0_seq1; Table 3; Figure 2a; 15 °C to 5 °C) and −1.39 (comp6405_c0_seq1; Table 4; Figure 3a; 5 °C to 0 °C), respectively. Yop1 is a membrane protein that associates with several other proteins to form the protein transport machinery of eukaryotic cells. More specifically, Yop1 was reported to be associated with another protein known as Yip1p which is essential for mediating membrane traffic (Calero et al. 2001). These proteins regulate many steps of the membrane traffic, including vesicle formation, vesicle movement along the actin and tubulin networks, and membrane fusion. They are required to undergo a cycle of binding and hydrolysis during protein transportation (Calero et al. 2001). The reduction of *YOP1* expression could be an effect of the decrease in cell membrane fluidity and membrane-associated functions due to the cold. In general, the rigidification of the membrane normally acts as the primary signal observed by psychrophilic microorganisms when exposed to low temperature before a series of cold adaptation mechanisms take place (Shivaji and Prakash 2010). The down-regulation of *YOP1* in USM-PSY62 in response to the cold-shift could possibly be to avoid cell death. Conversely, an over-expression of *YOP1* could produce an adverse effect by inhibiting growth and enlarging cellular size (Calero et al. 2001). Cellular enlargement was attributed to an accumulation of internal membrane structures. An over-expression of *YOP1* could also block the transport and trafficking of vesicles from the endoplasmic reticulum to the Golgi apparatus (Calero et al. 2001).

The expression of the gene for murein lipoprotein (comp12847_c0_seq1; Table 3; Figure 2a) was up-regulated by log2 fold change values of 9.71 during the cold-shifting condition of 15 °C to 5 °C. This could be a response to restore the fluidity of cell membrane. The murein lipoprotein is one of the components involved in the outer cell membrane envelope biogenesis. Several reports have seen that the murein lipoprotein is involved in the assembly of cell envelope (Braun and Bosch 1972), cell division (Torti and Park 1976; Weigand et al. 1976), and passive transport of small molecules through the outer membrane (Inouye 1974). The up-regulation of the murein lipoprotein gene could also be a compensating response for the reduced expression of *YOP1* which had decreased the cell membrane fluidity and membrane-associated functions.

The expression of the gene responsible for the structural maintenance of chromosome (*SMC*) showed a down-regulation with a log2 fold change value of −8.80 (comp6918_c0_seq1; Table 4; Figure 3a) when the temperature was changed from 5 °C to 0 °C. This family of SMC proteins are important in the proper segregation of chromosomes during mitosis and a wide variety of chromosomal events (Hirano 2006) (Losada and Hirano 2005; Nasmyth and Haering 2005). They are present abundantly in almost all organisms from bacteria to human (Hirano 2006). They are normally involved in chromosomal condensation, segregation, cohesion, and recombinational DNA repair in eukaryotes as well as introducing positive supercoiling into DNA *in vitro* (Graumann 2001). The result of the down-regulation of the genes for SMC protein in USM-PSY62 implicated a change in chromatin structure as a cold shock response. This observation is in accordance to a previous study which showed that the curvature of DNA changed and tended to become more negatively supercoiled at low temperatures (Barria et al. 2013). The down-regulation of SMC proteins conceivably resulted in the negative supercoiling of nucleic acids as a response towards cold temperature. The increased DNA negative supercoiling might consequently elicit more cold adaptive responses (Barria et al. 2013).

At cold temperatures, both transcriptional and translational activities of cells are reduced due to increased transient stabilization of DNA and RNA (Satyanarayana et al. 2005). To allow continuity of protein expression, cells tend to over-produce DNA and RNA helicases to reorganize the DNA and RNA secondary structures (Barria et al. 2013). This adaptive response was also observed in USM-PSY62 in which a putative homologue for DNA/RNA helicase, HepA, was over-expressed in cold condition. The *hepA* homologues were up-regulated the most at log2 fold change values of 2.89 (comp8191_c0_seq3; Table 3; Figure 2b; 15 °C to 5 °C) and 6.03 (comp8191_c0_seq3; Table 4; Figure 3b; 5 °C to 0 °C), as well as 9.04 (comp8936_c0_seq12; Table 3; Figure 2b; 15 °C to 5 °C) and 10.2 (comp8936_c0_seq8; Table 4; Figure 3b; 5 °C to 0 °C), respectively. This observation is in accordance with the results reported by a previous study on psychrophiles (De Maayer et al. 2014). The *hepA* homologue was one of the many genes that are strongly up-regulated in response to cold exposure. Cold-induced RNA helicases can destabilize secondary DNA and RNA structures and prevent possible interference of transcription (De Maayer et al. 2014).

Cellular metabolisms are important in providing the necessary components and energy to drive the cellular machineries for survival. At low temperatures, active Antarctic basidiomycetous yeast cells were seemingly focused towards metabolisms for survival rather than growth (Amato et al. 2009). In USM-PSY62, the expression of the homologue coding for CorA was up-regulated with a log2 fold change value of 7.0 (comp8278_c0_seq4; Table 3; Figure 2c; 15 °C to 5 °C) under cold-shock condition. The CorA protein family forms an important group of magnesium transporters in both prokaryotic and eukaryotic cells. The proteins function in facilitating magnesium uptake and distributing the magnesium ions to the subcellular compartments (Moomaw and Maguire 2008). The enhanced transcription of the homologues for CorA in USM-PSY62 showed an increased requirement for magnesium ions as cofactors of the enzymes and proteins induced during the cold-shock conditions. The CorA proteins do not necessarily act solely as magnesium ion transporters but possibly as other metal ion transporters as well. In prokaryotes, the CorA proteins are also involved in the uptake of cobalt ions (Hmiel et al. 1986). There is also a possibility that the increased presence of the CorA proteins resulted in an increased ionic concentration within the cell cytoplasm. A high concentration of ions decreases the freezing point of water or cytoplasmic fluid, thus preventing ice crystal damages which can be fatal to cells (Sherwood et al. 2012). In order to survive, USM-PSY62 could have increased its ionic intake to maintain the fluidity of its cytoplasm. Different strategies to maintain cell fluidity have been observed in other Antarctic species of yeast. *Mrakia psychrophila* and *Mrakia blollopis* up-regulated their glycerol 3-phosphate dehydrogenase genes to prevent osmotic shrinkage during cold stress acclimation (M 2018; Su Y 2016; Wolfe J 1999). The sea ice psychrophilic basidiomycete *Glaciozyma antarctica* PI12 was equipped with nine antifreeze proteins to adapt to the sub-zero freezing conditions (Firdaus-Raih M 2018).

Another common physiological response of organisms to different types of environmental stresses is the increase in expression of heat shock proteins (HSPs). Heat shock genes are induced as an adaptive response towards shifts to high temperature conditions. A rapid increase in environmental temperature can result in severe damage to cellular proteins and membranes and most of the time, proteins lose their conformational flexibility leading to protein malfunction (Richter et al. 2010). Thus, HSPs provide protection by acting as molecular chaperones, thereby alleviating the harmful effects of misfolding and degradations. HSPs have been widely studied for heat stress but the response to cold stress has been far less investigated. HSPs are actually associated with both extremely cold- as well as heat-shock conditions. During periods of stress, HSPs are rapidly upregulated as a response until when preferable conditions prevail again (Rinehart 2007).

Although various reports have described the heat shock response of mesophilic and thermophilic microorganisms, little is known about the phenomenon in psychrophilic microorganisms (Yumoto 2013). Psychrophilic yeasts are able to synthesize heat shock proteins (HSPs) as a defence mechanism (Buzzini et al. 2012). In this study, when the temperature shifted down, the gene for heat shock factor protein 1 (HSF1) was down-regulated as a response (comp6218_c0_seq1; Table 4; Figure 3b; 15 °C to 5°C, comp5468_c0_seq1; Table 4; Figure 3b; 5 °C to 0 °C, and comp5468_c0_seq1; Table 3; Figure 2b; 15 °C to 5 °C). The Hsf1 protein belongs to a group of transcriptional regulators known as heat shock factors (HSFs) which regulate the heat shock responses. Under normal condition, Hsf1 exists in a complex with either Hsp40/70 or Hsp90 as an inactive monomer (Branda et al. 2010). When the cells are under thermal stress, Hsf1 is released from the chaperone complex and undergo trimerization. Hsf1 is subsequently transported into the nucleus, hyperphosphorylated and eventually binds to the DNA containing heat shock elements consisting of the repeating five-nucleotide units of NGAAN (Shamovsky and Nudler 2008). Hsf1 has a helix-turn-helix DNA-binding domain to enable the attachment to occur (Bonner 1992). Interestingly, over-expression of *HSF1* arrest the vegetative growth of haploid yeast cells while a deletion or a null mutant is lethal. Temperature sensitive mutants are defective in several processes including preservation of cell wall integrity, duplication of spindle pole body, transportation of proteins (Boeke 1984; Mosser 1990; Oettinger 1985; Sadler 1989).

Up-regulation of the gene for catalase (CAT, EC 1.11.1.6) was one of the distinct responses of USM-PSY62 during the stepped down-shifts of temperature. Catalase is an antioxidant enzyme existing in all aerobic organisms. It decomposes hydrogen peroxide (H2O2) to water and molecular oxygen. At low H2O2 concentrations, the enzyme oxidizes low molecular alcohols, nitrites and formate that are generated during biological oxidation (Kurutas 2016). The expression of CAT in USM-PSY62 was up-regulated with log2 fold change value of 2.6 (comp2593_c0; Table 3; Figure 2c; 15 °C to 5 °C) when temperature shifted from 15 °C to 5 °C. A similar observation was made in *Saccharomyces cerevisiae* in which a downshift in the growth temperature from 30 °C to 10 °C resulted in an increase in transcript levels of the genes encoding catalase T (CTT1), Cu–Zn superoxide dismutase (SOD1) and γ-glutamylcysteine synthetase (GSH1) (Zhang 2003). These are genes involved in antioxidant response indicating that down-shift in temperature could lead to oxidative stress. In *Arabidopsis thaliana* and wheat, low temperature stress thrusted the sub-cellular antioxidant systems to protect photosynthetic electron transport (Caverzan 2016; Juszczak 2016).

Metabolic strategies to overcome low-temperature conditions are also involved in cold adaptation. For example, genes involved in carbon metabolism and energy production were found to be differentially expressed in *Glaciozyma antarctica* PI12 and *Mrakia blollopis* (Firdaus-Raih M 2018; M 2018). The same physiological strategies were utilized by USM-PSY62 to compensate for the low-temperature effect. For example, homologues of ProP (permeases of the major facilitator superfamily; comp2909_c1_seq1; Table 4; Figure 3c), BglX (beta-glucosidase-related glycosidases; comp8366_c0_seq2 & comp8366_c0_seq4; Table 4; Figure 3c), and LacZ (beta-galactosidase; comp3495_c0_seq2; Table 4; Figure 3c) involved in carbohydrate transport metabolism were significantly upregulated at the lower temperatures. Genes involved in amino acid transport and metabolism such as those coding for PotE (amino acid transporter; comp8592_c0_seq12; Table 4; Figure 3c) and BetA (choline dehydrogenase and related flavoproteins; comp9040_c0_seq32; Table 4; Figure 3c) were also upregulated. A gene involved in energy production and conversion, namely *qor*, a NADPH:quinone reductase and related Zn-dependent oxidoreductase (comp8089_c0_seq4, comp8089_c0_seq3, comp8089_c0_seq2; Table 4; Figure 3c) were also transcriptionally enhanced.

In conclusion, the transcriptomic responses demonstrated by USM-PSY62 characterized critical physiological and biochemical compensatory mechanisms to survive at low as well as high temperatures. The responses have relevance to its survival and are important for the endurance of eukaryotic microbes in the freezing Antarctic environment. Bioinformatics studies on various species other than model organisms are crucial to uncover new molecular pathways or new genes responsible for adaptation in thermal stresses.

## Supporting information

ESM 1

## Acknowledgements

This research was funded by the Malaysian Ministry of Energy, Science, Technology and Climate Change (MESTECC) under the Genomics and Molecular Biology Initiatives (Science Fund Project No: 08-05-MGI-GMB001). We acknowledge support given by the Australian Antarctic Division, Sultan Mizan Antarctic Research Foundation and the Malaysian Antarctic Research Program (MARP) of the Academy of Science, Malaysia. Authors declare no conflict of interest.

## Notes

https://www.ncbi.nlm.nih.gov/bioproject/PRJNA522043

### Competing Interest Statement

The authors have declared no competing interest.

