## Supplementary material for "Transcriptomic response of an Antarctic yeast *Rhodotorula* sp. USM-PSY62 to temperature changes": ESM 1

**Title**

[3] Current address: Faculty of Applied Science, University College Sedaya International (UCSI), 56000 Kuala Lumpur, Malaysia

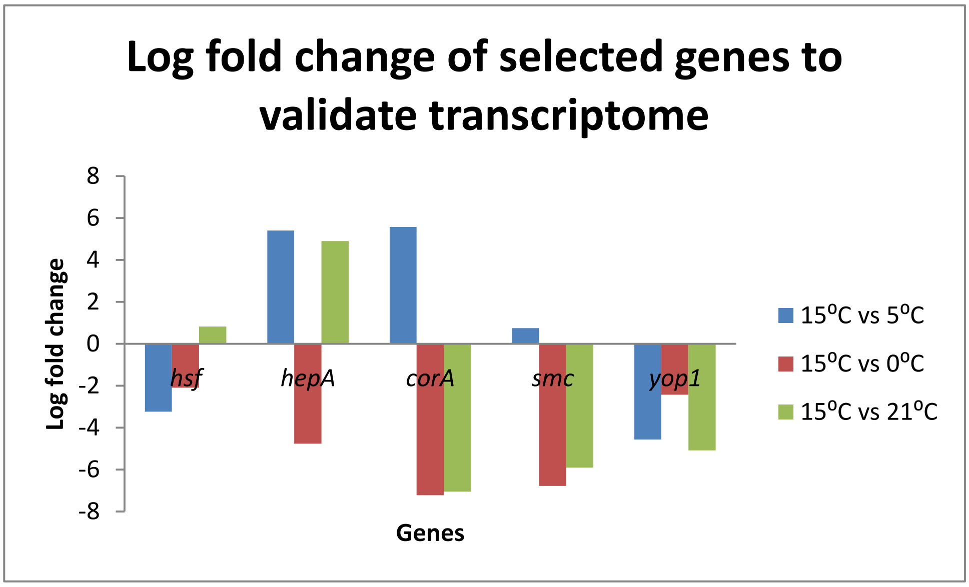


**Online Resource 1 Log fold change of *hsf*, *hepA*, *corA*, *smc* and *yop1* genes.** The validation of transcriptome analysis was performed using qRT-PCR. The relative expression of each genes were normalized using the expression of control gene, actin and analyzed using Livak's ($2^{-\Delta\Delta Cq}$) method. The result corresponded with transcriptome analysis and thus validated bioinformatics analysis.
